# Perceptron learning and classification in a modeled cortical pyramidal cell

**DOI:** 10.1101/464826

**Authors:** Toviah Moldwin, Idan Segev

## Abstract

The perceptron learning algorithm and its multiple-layer extension, the backpropagation algorithm, are the foundations of the present-day machine learning revolution. However, these algorithms utilize a highly simplified mathematical abstraction of a neuron; it is not clear to what extent real biophysical neurons with morphologically-extended nonlinear dendritic trees and conductance-based synapses could realize perceptron-like learning. Here we implemented the perceptron learning algorithm in a realistic biophysical model of a layer 5 cortical pyramidal cell. We tested this biophysical perceptron (BP) on a memorization task, where it needs to correctly binarily classify 100, 1000, or 2000 patterns, and a generalization task, where it should discriminate between two “noisy” patterns. We show that the BP performs these tasks with an accuracy comparable to that of the original perceptron, though the memorization capacity of the apical tuft is somewhat limited. We concluded that cortical pyramidal neurons can act as powerful classification devices.

## Introduction

There has been a long-standing debate within the neuroscience community about the existence of ‘grandmother neurons’ — individual cells that code for high-level concepts — such as a person’s grandmother. Recent experimental evidence, however, has indicated that there are units that are selective to specific high-level inputs. In particular (Quiroga *et al.*, 2005) found cells in the human medial temporal lobe (MTL) that fire in response to images of a particular celebrity, such as Jennifer Aniston or Halle Berry. One remarkable aspect of this finding is that different images of the same celebrity would elicit a response in these neurons even if the subject of the image was facing a different direction, wearing different clothes, or under different lighting conditions. In other words, the specificity of these MTL cells is invariant to certain transformations of the sensory stimulus. Regardless of whether this finding is evidence for grandmother cells or merely for sparse coding (Quiroga *et al.*, 2008), it is apparent that individual neurons can be highly selective for a particular pattern of sensory input and also possess a certain level of generalization ability, or “tolerance”, to differences in the input that do not change the essence of the sensory scene.

From a physiological standpoint, achieving a high degree of accuracy on a recognition task is a daunting challenge for a single neuron. To put this in concrete terms, a pyramidal neuron may receive around 30,000 excitatory synapses (Megías *et al.*, 2001). As a first approximation, at any given moment, each of this neuron’s presynaptic inputs can either be active or inactive, yielding 2^30,000^ possible binary patterns. If the presynaptic inputs contain information about low-level sensory stimuli (such as pixels or orientation filters) and the postsynaptic neuron needs to respond only to images of Jennifer Aniston, for example, there must be some physiological decision procedure by which the neuron “chooses” which of those 2^30,000^ patterns are sufficiently close to the binary representation of Jennifer Aniston to warrant firing a spike as output.

There are several ways that a neuron can selectively respond to different input patterns. The most well-known method is to adjust synaptic “weights” such that only input patterns which activate a sufficient number of highly-weighted synapses will cause the cell to fire. It is this principle which serves as the basis of the perceptron learning rule (Rosenblatt, 1958) which is, in turn, the foundation for the artificial neural networks (ANNs) that are commonly used today in machine learning and deep networks (Rumelhart, Hinton and Williams, 1986; Krizhevsky, Sutskever and Hinton, 2012).

The perceptron utilizes a mathematical abstraction of a neuron which applies a nonlinear function (such as a sigmoid) to the weighted sum of its input (Figure 1A). This abstraction is known as the McCulloch and Pitts (M&P) neuron (McCulloch and Pitts, 1943). The nonlinear output of the neuron plays the role of a classifier by producing a positive output (a spike, +1) in response to some input patterns and a negative output (no spike, −1) in response to other patterns. This “perceptron” neuron model is trained in a supervised manner wherein it receives training patterns which are labeled as belonging to either the positive or negative category. The perceptron output is calculated for each pattern, and if the perceptron output for a particular pattern does not match the label, the perceptron’s weights are updated such that its output will be closer to the correct output for that example in the future.

**Figure 1.**
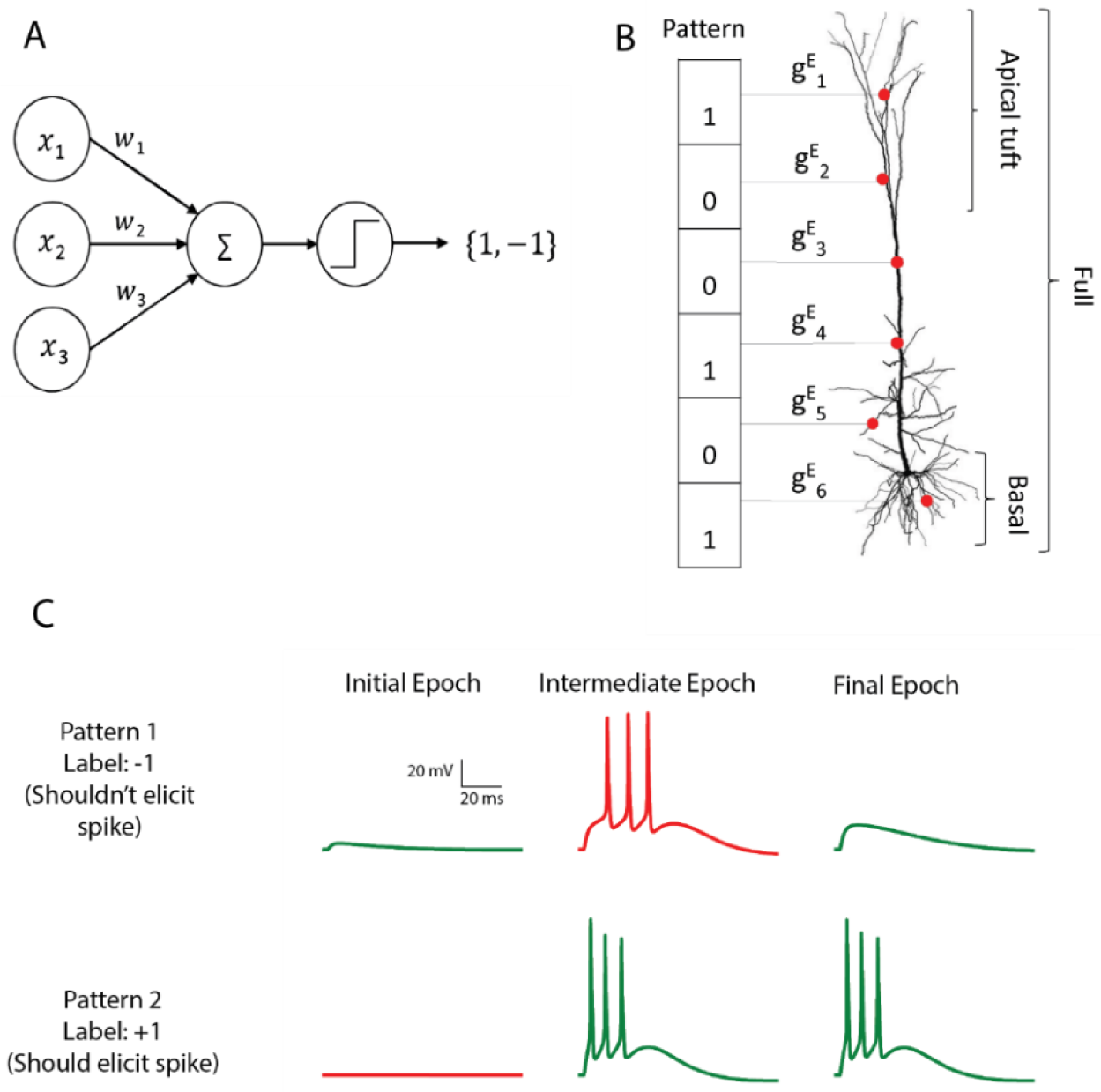
The M&P and biophysical perceptron. (A) The M&P perceptron. The presynaptic neurons are represented by their firing rates, *x*_1_, *x*_2_, *x*_3_, … x_i_, each of which is multiplied by the respective synaptic weight *w*_1_, *w*_2_, *w*_3_, … *w*_*i*_ and then summed together with the other inputs. The perceptron produces an output of +1 if the weighted sum of all inputs is greater than a threshold and −1 otherwise. The task of the perceptron is to learn the appropriate synaptic weights, such that it will produce an output of +1 for an arbitrarily predefine subset of *x*_*i*_, and −1 for the remaining subset of *x*_*i*_. (B) Schematic of the biophysical perceptron. A layer 5 pyramidal cell model with excitatory synapses (red dots) receiving an exemplar presynaptic input pattern. The synaptic weights are the excitatory conductance, *g*^*E*^ _*i*_, for the respective synapse, *i*. In this model, a presynaptic input pattern consists of a particular set of synaptic inputs that are either active “1” or inactive “0”. (C) An example of the learning process in the biophysical perceptron. Two input patterns, each with 1000 synapses, were presented to the model neuron. For pattern 1 the model cell should not generate any spike, whereas for pattern 2 it should. In the initial epoch neither pattern elicits a spike (traces at left). The output for the pattern 1 is thus correct (green trace) but incorrect (red trace) for pattern 2. In an intermediate epoch of the learning algorithm (middle column), some of the synaptic conductances were sufficiently increased so that pattern 2 does elicit spikes, however pattern 1 also produces spikes (but it should not). By the final epoch (left column), the weights are adjusted such that the neuron correctly classifies the two patterns (right column).

While the remarkable efficacy of networks of M&P neurons has demonstrated for various learning tasks, few attempts have been made to replicate the perceptron learning algorithm in a detailed biophysical neuron model with a full morphology and active dendrites with conductance-based synapses. It thus remains to be determined whether real cells in the brain, with all their biological complexity, can integrate and classify their inputs in a perceptron-like manner.

In this study, we used the perceptron learning algorithm to teach a detailed realistic biophysical model of a layer 5 pyramidal cell (Hay et al., 2011) to solve two kinds of classification problems: a memorization task, where the neuron must correctly classify (by either spiking or not) a predefined set of ‘positive’ and ‘negative’ input patterns, and a generalization task, in which the neuron has to discriminate between two patterns that are corrupted by noise in the form of bit flips (i.e. where active synaptic inputs are switched to inactive and vice versa). We explored the ability of real neurons with extended nonlinear dendritic trees and conductance-based excitatory synapses to perform classification tasks of the sort commonly solved by artificial neurons (see **Discussion** for a treatment of why only excitatory synapses were used). We found that the performance of the biophysical perceptron (BP) is close to that of the artificial M&P counterpart.

## Results

### Memorization task

To implement the perceptron learning algorithm in a modeled layer 5 thick tufted pyramidal cell (L5PC) we distributed conductance-based excitatory, AMPA and NMDA-based, synapses on the detailed model developed by (Hay *et al.*, 2011). We created input patterns consisting of 1000 excitatory synapses, 200 of which were active in any given pattern. We varied the total number of patterns (P) presented to the modelled neuron in order to determine its classification capacity (Figure 1B). We tested conditions of P = 100, P = 1000, and P = 2000. These binary patterns were evenly divided into a “positive” (+1) group (for which the modeled neuron should produce at least one spike) and a “negative” (−1) group (for which the modeled neuron should not produce a spike). To achieve perfect accuracy, the neuron would have to correctly fire in response to all the patterns in the positive group and not fire in response to all the patterns in the negative group. Note that, initially, there is no reason for the neuron to perform at better than chance level, because all the patterns contain the same number of active synapses.

We then used the perceptron learning algorithm (see **Experimental Procedures**) to modify the synaptic weights such that the cell could correctly classify all the patterns (Figure 1C). This procedure was repeated in conditions in which synapses were placed over the whole dendritic tree, only on the apical tuft, only on the basal tree, or only on the soma in order to determine how the location of the synapses affects the cell’s ability to classify patterns using the perceptron learning rule (see **Discussion** for the biological significance of input patterns on different parts of the dendritic tree). We also tested the algorithm with current-based synapses rather than of conductance-based synapses, to examine whether conductance-based synapses have any advantages or disadvantages with respect to the cell’s performance as a classifier.

Figure 2 shows the learning curves (Figure 2A) and classification accuracy (Figure 2B) for each of the above-mentioned conditions. In all cases the cell is able to improve its performance relative to chance, indicating that the complexity of biophysical cells does not preclude perceptron learning despite the fact that the learning algorithm having been devised for a much simpler abstraction of a cell.

**Figure 2.**
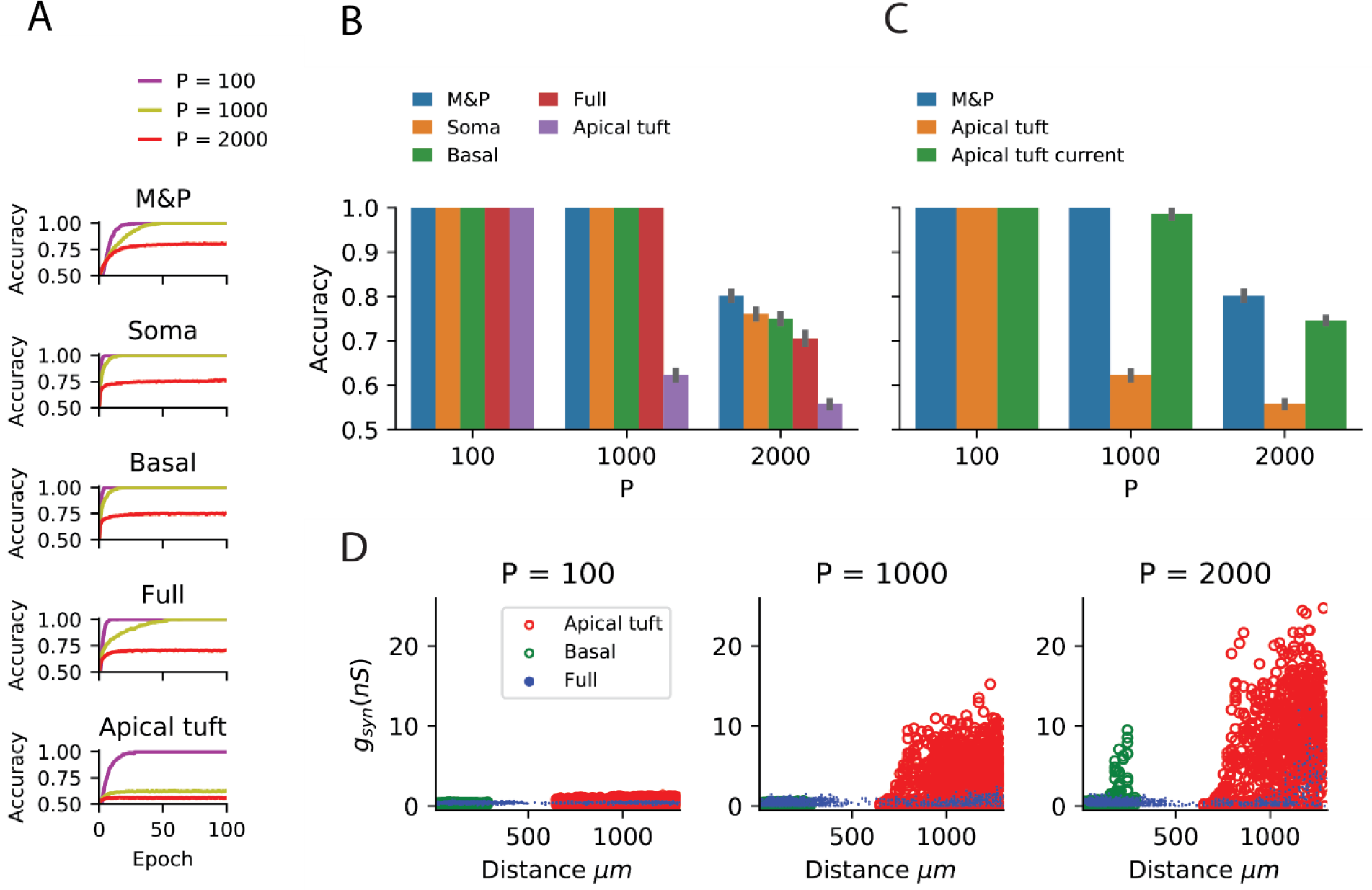
Learning the memorization task with the biophysical perceptron. (A) Learning curves for the memorization task, in which a neuron with 1000 excitatory synapses had to classify patterns (half of which should produce a spike, half should not). The results for tasks involving 100, 1000 and 2000 patterns are shown (P, colored traces). The learning curves of an M&P perceptron is depicted at the top and the four different conditions of synaptic placement in the L5PC model shown in Figure 1 are shown subsequently. **Soma**: all synapses are placed only at the modeled soma; **Full**: all synapses are uniformly distributed (per unit length) on the whole dendritic tree; **Basal**: all synapses are uniformly distributed on the basal tree (see Figure 1B); **Apical tuft**: all synapses are uniformly distributed on the apical tuft (see Figure 1B). (B) Accuracy in the memorization task for different synaptic placement conditions. Mean accuracy after 100 epochs is shown (error bars: standard deviation) for different numbers of patterns (P). The blue bar within each grouping shows the performance of a M&P neuron. Note the poor performance in the apical tuft placement condition. (C) Effect of conductance synapses vs. current synapses on classification capacity in the apical tuft. As in (B), blue bar is the performance of an M&P neuron for comparison. Note that classification capacity of the apical tuft is restored to near that of the M&P perceptron when switching from conductance synapses to current synapses. Error bars as in (B). (D) Value of synaptic conductances obtained after the completion of the learning algorithm as a function of distance of the synapses from the soma for learning tasks with different numbers of patterns. The cases of synapses placed only on the apical tuft (red), basal tree (green) and full (blue) placement conditions are shown for a single exemplar run. Note the large synaptic conductances obtained during the learning task for the case where the synapses are placed at the apical tuft for P = 1000 and 2000.

We compared the classification accuracy for each condition in the biophysical model to an equivalent M&P perceptron with excitatory weights (see **Experimental Procedures**). When all synapses are placed on the soma or the proximal basal tree of the biophysical perceptron, the classification accuracy of the biophysical perceptron is near to that of the M&P perceptron.

As expected from the theoretical literature (Julio Chapeton *et al.*, 2012), the accuracy in each condition decreases with the number of patterns that the neuron must learn. This can be seen in Figure 2B, where the classification accuracy degrades in each condition as we move from P = 100 to P = 1000 and from P = 1000 to P = 2000.

In all synaptic placement conditions, the M&P perceptron and the BP performed with perfect accuracy on the ‘easy’ task with P = 100. In conditions where the synapses were placed only on the soma or only on the basal tree, the performance of the BP is comparable to that of the M&P neuron for P = 1000 (M&P: 100%, basal: 100%, soma: 100%) and for P = 2000 (M&P: 80.3%, basal: 75%, soma: 76%). In the condition where synapses were placed uniformly over the full tree, the discrepancies were somewhat larger for P = 2000 (M&P: 80.3%, full: 70.5%).

However, when the synapses are all placed on the apical tuft of the biophysical cell, the classification accuracy of the biophysical perceptron decreases dramatically, even in the presence of supra-linear boosting mechanisms such as NMDA receptors and active Ca+ membrane ion channels. For P = 1000, the M&P neuron achieves 100% classification accuracy, whereas if the synapses are all placed on the apical tuft, the neuron only achieves 62% accuracy. In the condition with P = 2000, the M&P neuron achieves 80% classification accuracy whereas the BP achieves only 55.8% classification accuracy, barely better than chance level. However, by switching from conductance-based synapses to current-based synapses in the apical tuft condition, it was possible to regain almost all of the “loss” in the classification accuracy (In the P= 1000 condition, from 62% with conductance synapses to 98.5% with current synapses, in the P= 2000 condition, from 55.8% with conductance synapses to 74.5% with current synapses) (Figure 2C).

We argue that the reason for the discrepancy in classification accuracy for the biophysical perceptron between the conditions wherein synapses are placed on the apical tuft, as opposed to the soma or basal dendrites, is due to the passive filtering properties of the neuronal cable and the saturation effect of conductance synapses. Specifically, the attenuation of voltage along the length of cable from apical tuft dendrites to the spike initiation zone means that the *effective weight* of that synapse - namely the magnitude of the resultant somatic EPSP - is greatly reduced. This phenomenon has been observed previously (Rall, 1967; Stuart and Spruston, 1998), but it has been argued (Häusser, 2001; Rumsey and Abbott, 2006) that the cell might be able to overcome this drop in voltage by simply increasing the strength (i.e. conductance) of distal synapses. We demonstrated, however, that this is not the case. We show (Figure 2D) that the perceptron learning algorithm will, on its own, increase the weights of apical tuft synapses far beyond the biologically plausible range of 0.2 - 1.3 nS (Sarid *et al.*, 2007; Eyal *et al.*, 2018) in attempting to correctly classify all the patterns. Still, the classification accuracy of the apical tuft biophysical perceptron remains quite poor (see, however (Gidon and Segev, 2009) who show that the opposite phenomena will occur with a standard STDP rule, resulting in smaller synaptic conductances for distal synapses).

We claim that “democratization” via disproportionally increasing distal synaptic conductances does not solve the classification accuracy problem for synapses located on the apical tuft because effective synaptic weights are bounded by the synaptic reversal potential in the distal dendrites, even if one were to increase synaptic conductances to arbitrarily high values. As such, the maximal effective synaptic weight (MESW) - defined as the peak somatic EPSP voltage - when a given dendritic location attains the synaptic reversal potential (Figure 3A) - is equivalent to the synaptic driving force multiplied by the attenuation factor from that dendritic location to the soma. [Note: This is true in the passive case, dendritic nonlinearities can affect the MESW values. Our calculations of MESWs in this study are based on simulations of the model with all nonlinearities present, as shown Figure 3A.] The MESWs for distal synapses are thus smaller than those for proximal synapses (Figure 3B). Because the apical tuft is electrotonically distant from the soma, synapses on the apical tuft have lower MESWs (Median for apical tuft: 1.2 mV, full cell: 10.5 mV, basal tree: 12.15 mV mv, Figure 3C). The fact that switching the apical synapses from conductance-based to current-based substantially improves classification accuracy supports the notion that voltage saturation due to synaptic reversal potential is responsible for the reduced performance of the apical tuft synapses (Figure 2C).

**Figure 3.**
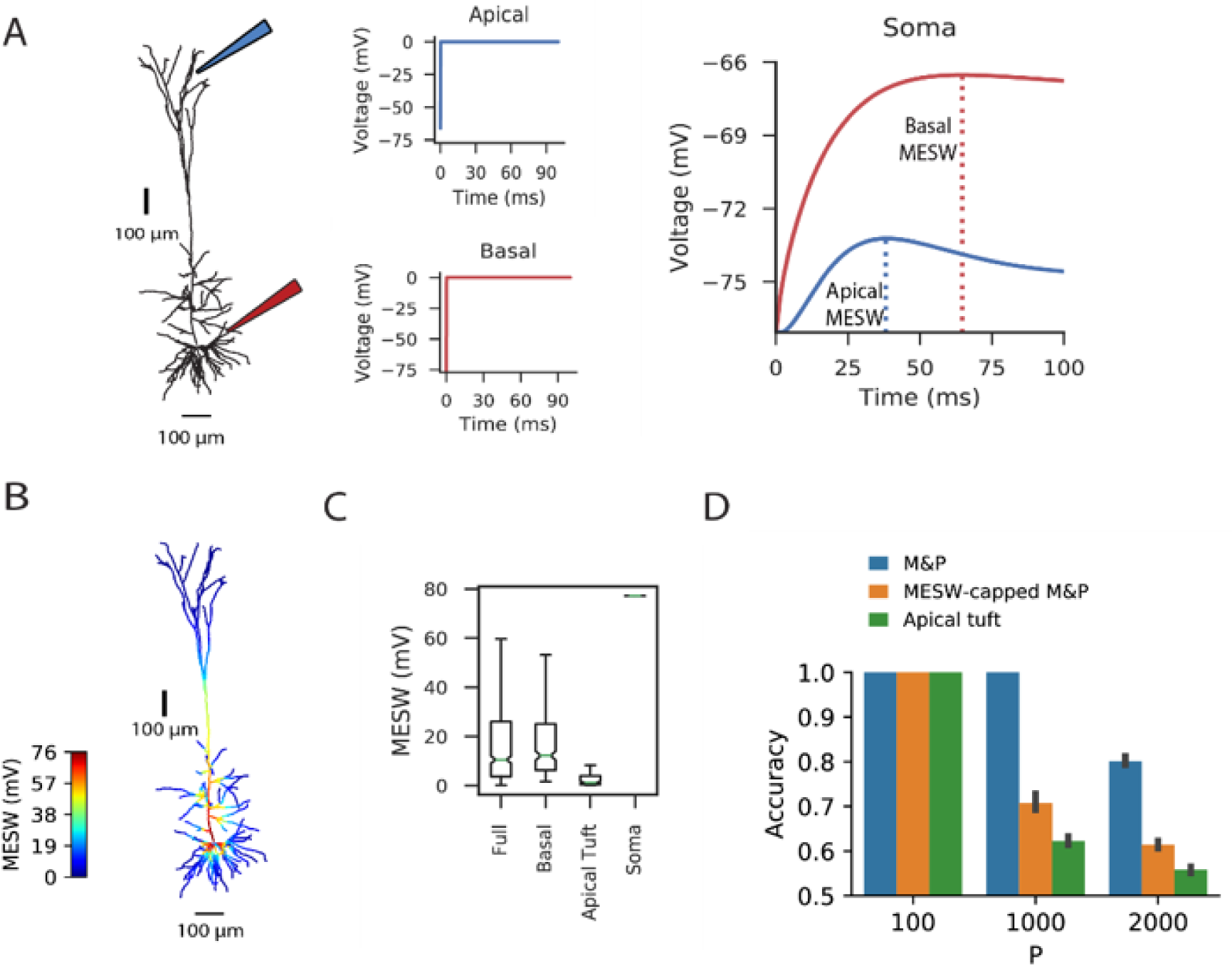
Effect of location of dendritic synapses on the maximum effective synaptic weight (MESW). (A) The simulated L5 pyramidal neuron with red (proximal basal) and blue (distal apical) electrodes. To calculate the MESW for each location, we voltage clamp the dendritic segment to 0 mV, the reversal potential for AMPA/NMDA synapses, for 100 ms (middle column) and record the resultant somatic voltage (right column). The MESW for two dendritic loci (dashed lines) is defined as the voltage difference between the somatic resting potential and the peak voltage observed at the soma during the dendritic voltage clamp. Note that the decline of the somatic depolarization after the peak is due to nonlinear dendritic ion channels. (B) A simulated L5 PC with the MESW values superimposed for each dendritic segment. Note the steep voltage attenuation from distal dendritic branches to the soma (blue regions). (C) Box-and-whiskers plot of MESW distributions for the full dendritic tree, basal tree, apical tuft, and soma. (see **Experimental Procedures)**. Notches represent the median MESWs for all synapses within that region, box edges and error bars respectively represent the first and second quartiles of the data. (D) Effect of constraining the synaptic weights in the M&P model according to the distribution of MESWs observed for the apical tuft. Note that the classification capacity of the MESW-constrained M&P perceptron is substantially reduced and becomes closer to the capacity of the biophysical perceptron.

From the standpoint of learning theory, the “cap” on the effective weights of distal apical synapses restricts the parameter space of the biophysical perceptron, reducing its capacity. When a perceptron learns to classify between two sets of patterns, it creates a linear separation boundary – i.e. a hyperplane – which separates the patterns in an N-dimensional space, where N is the number of inputs\synapses in each pattern. The separation boundary learned by the perceptron is defined by the hyperplane orthogonal to the vector comprising the perceptron’s weights. When the weights of the perceptron are unconstrained, the perceptron can implement any possible hyperplane in the N-dimensional space. However, when the weights are constrained – for example by the MESWs of the apical tuft of L5 PCs – the perceptron can no longer learn every conceivable linear separation boundary, reducing the ability of the perceptron to discriminate between large numbers of patterns. [Note: because we use only excitatory synapses, the weight space in all synaptic placement conditions is already substantially constrained to positive values even before imposing MESWs, see (J. Chapeton *et al.*, 2012) for a full treatment.] To demonstrate this effect, we calculated the MESW for each synapse in the apical tuft and then imposed this distribution of MESWs onto an M&P perceptron (see **Experimental Procedures**). As predicted, the MESW-capped M&P perceptron produced a reduced classification capacity in a manner similar to the biophysical perceptron when synapses were restricted to the apical tuft (Figure 3D).

It should be noted that the limited capacity of the apical tuft is *not* because apical synapses cannot induce the neuron to fire, as the neuron with only apical synapses performs with perfect accuracy when it only needs to classify 100 patterns, indicating that 200 active synapses on the apical tuft are fully capable of generating a somatic spike. It is thus evident that the small classification capacity of the apical patterns is due to the restriction of the weight space needed to properly discriminate between positive and negative patterns, not because the apical tuft input is insufficiently strong to create a somatic spike.

### Generalization task

To explore whether the apical tuft is always at a disadvantage when it comes to pattern classification, we also tested the biophysical perceptron on a generalization task. Instead of “memorizing” a large set of fixed patterns, in the generalization task the neuron was presented with “noisy” patterns drawn from one of two underlying fixed patterns. In this task, noise was added to the underlying pattern by performing “bit flips”, i.e. flipping an active synapse to an inactive synapse or vice versa (Figure 4A). We tested both the biophysical perceptron (with different synaptic placement conditions, as in the memorization task) and the positive-weighted M&P neurons on their ability to classify these noisy patterns in conditions with varying levels of difficulty, as determined by the number of bit flips. The goal of the task was that the neuron should fire in response to noisy patterns generated by the first underlying pattern, but not fire in response to noisy patterns generated by the second underlying pattern (Figure 4B).

**Figure 4.**
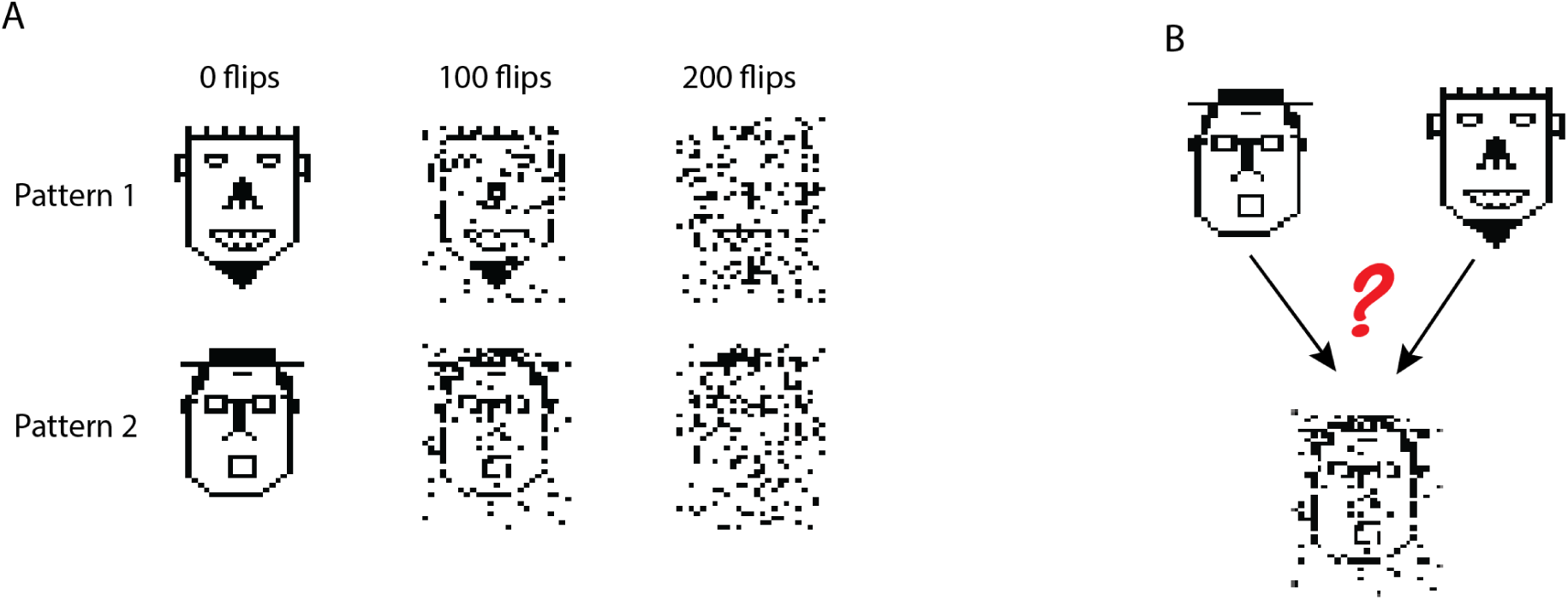
Generalization task with the biophysical perceptron. (A) Left column: Two binary patterns, represented by two faces, each consisting of 200 black pixels and 800 white pixels. The black pixels represent active synapses and the white pixels represent inactive synapses. These patterns can be corrupted by flipping active synapses to inactive synapses or *vice versa*, creating a new “noisy” pattern (middle and right columns). As we increase the number of flipped synapses, the noisy patterns become more difficult to identify with the original pattern. The faces are for illustration of the sparsity and noise level only, actual input patterns were not created with a facial structure. (B) Task schematic. A noisy pattern (lower face), drawn from one of two original patterns (top faces) is presented to the modeled neuron. The neuron must decide from which of the two original patterns the noisy pattern came, by either firing (for the first pattern) or not firing (for the second pattern).

In this task, we observe that in all conditions the BP performs similarly to the M&P perceptron. We do not observe any substantial diminution in classification performance between the apical tuft and the soma, as we do in the memorization task (Figure 5A-5B). In the condition with 100 bit flips, the difference in accuracy between the apical tree and the soma were small (M&P: 84% soma: 85%, apical tuft: 81.8%). The same is true for difficult task with 200 bit flips (M&P: 70.35%, soma: 71.8%, apical tuft: 67.4%). Changing the conductance synapses to current synapses did not substantially affect these results (Figure 5C). Moreover, capping the weights of the M&P neuron with the MESWs from the apical tuft, as we did in the memorization task, did not considerably worsen the M&P perceptron’s performance (Figure 5D). We also note that, while the some of the synaptic weights of the apical tuft did increase beyond the biological range during learning in the biophysical perceptron (Figure 5E), the effect is much smaller in the memorization task (Figure 2D).

**Figure 5.**
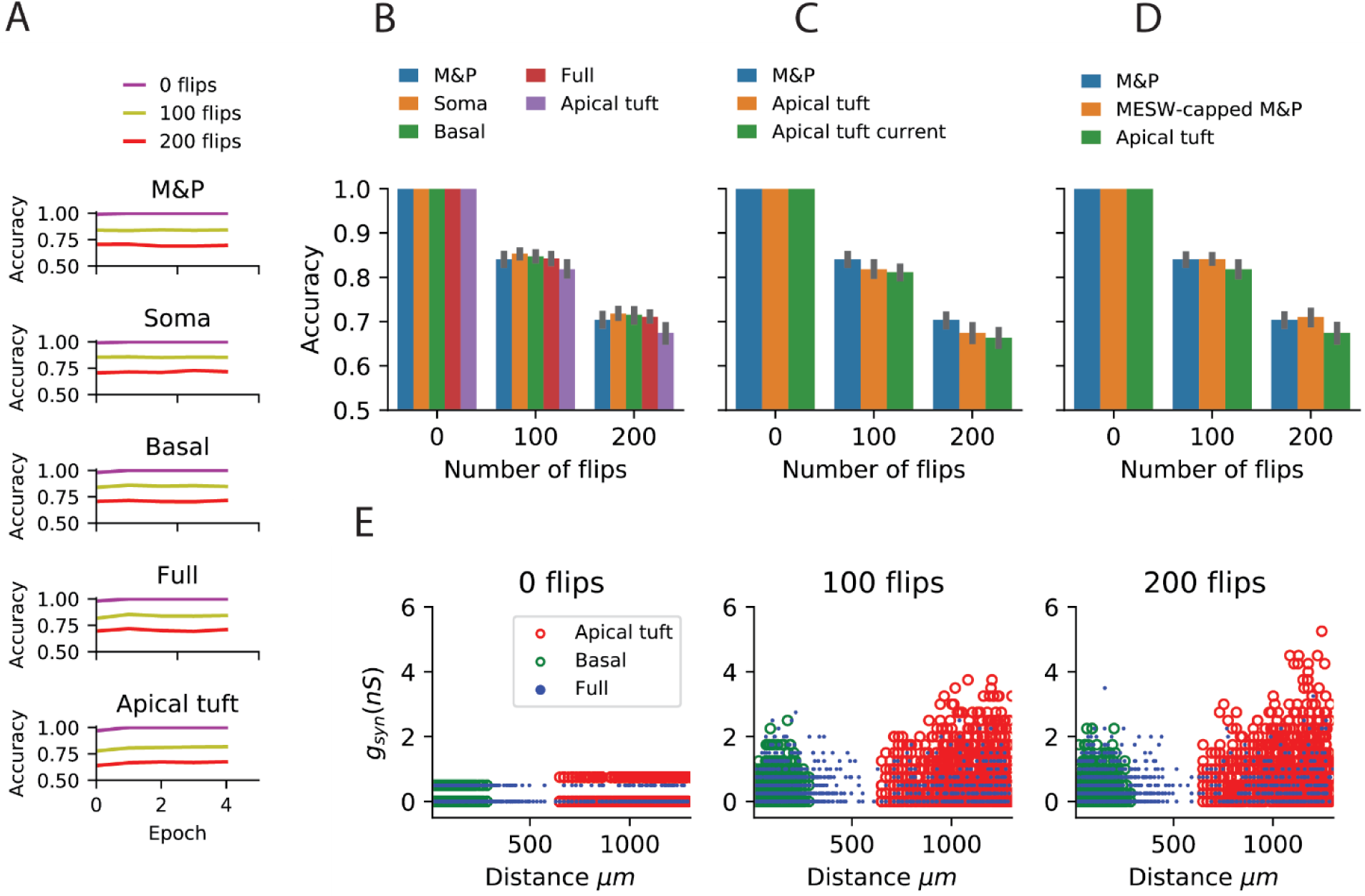
Learning the generalization task with the biophysical perceptron. (A) Learning curves for the generalization task, in which a neuron with 1000 synapses had to classify noisy patterns (see Figure 4) drawn from one of two underlying original patterns (see Figure 4). The neuron was presented with 100 different noisy patterns each epoch, 50 from each original pattern. The results for tasks involving different amounts of noise are shown (bit flips, colored traces). The learning traces of an M&P perceptron is depicted at the top and the four different conditions of synaptic placement in the L5PC model are shown subsequently. Synaptic placement as in Figure 2A. (B) Accuracy in the generalization task for different synaptic placement conditions. Mean accuracy after 100 epochs is shown (error bars: standard deviation) for different amounts of noise (bit flips). The blue bar within each grouping shows the performance of a M&P neuron. (C) Effect of conductance synapses vs. current synapses on classification capacity in the apical tuft. As in (B), blue bar is the performance of an M&P neuron for comparison. Error bars as in (B). (D) Effect of constraining the synaptic weights in the M&P model according to the distribution of MESWs observed for the apical tuft. (E) Value of synaptic conductances obtained after the completion of the learning algorithm as a function of distance of the synapses from the soma for learning tasks with different numbers of patterns. The cases of synapses placed only on the apical tuft (red), basal tree (green) and full (blue) placement conditions are shown for a single exemplar run.

The discrepancy between the apical tuft and soma may be smaller in the generalization task than in the memorization task because the difficulty in the memorization task is fundamentally about finding the correct hyperplane that will separate between the two classes of patterns. As we increase the number of patterns in each of the classes, we require more flexibility in the weight space of the neuron to ensure that all the positive and negative patterns end up on opposite sides of the separating hyperplane. This flexibility is impeded by the MESWs of the apical tuft. By contrast, the generalization problem only contains two canonical “patterns”. The difficulty in learning the generalization task with a large amount of noise (in terms of bit flips) does not stem from the challenge of precisely defining a separation boundary. Rather, solving the generalization task is hard because, even if we had an optimal separation boundary, the noise in the input entails that some of the noisy patterns would still necessarily be misclassified.

## Discussion

In the simulations described above, we have demonstrated that the perceptron learning algorithm can indeed be implemented in a detailed biophysical model of L5 pyramidal cell with conductance-based synapses and active dendrites. This is despite the fact that the perceptron learning algorithm traditionally assumes a cell which integrates its inputs linearly, which is not the case for detailed biophysical neurons with a variety of nonlinear active and passive properties and conductance-based synapses. Evidently, although such nonlinearities are present, they do not qualitatively prevent the cell from learning in a perceptron-like manner. That being said, the ability of a biophysical perceptron to distinguish between different patterns of excitatory synaptic input does depend on the location of the relevant synapses. Specifically, if all the synapses are located proximally to the soma, such as on the proximal basal tree, the cell has a classification capacity similar to that of the M&P perceptron. However, for activation patterns consisting of more distal synapses, such as those on the apical tuft, the classification capacity of the BP is reduced. We showed that this is due to the reduced effectiveness of distal synapses due to cable filtering and synaptic saturation, which limits the parameter space of the learning algorithm and thus hampers classification capacity. We also demonstrated that the diminished classification capacity in the apical tuft is negligible in a generalization task. This indicates that, while the maximum effective synaptic weights of the apical tuft may be somewhat limiting for its classification capacity, they do not hamper the apical tuft’s robustness to noise (Figure 5).

The above discussion considers that the pyramidal cell separately classifies inputs that synapse onto different regions of its dendrites (such as the apical tuft and the basal tree) and that it does not simultaneously integrate all the synaptic input impinging on the cell. This decision was motivated by a growing evidence that different parts of the dendritic tree may play separate roles in shaping the neuron’s output. From anatomical studies, it is known that axons from different brain regions preferentially synapse onto particular regions of layer 5 pyramidal cells. For example, basal dendrites tend to receive local inputs whereas the apical tuft receives long-range cortical inputs (Crick and Asanuma, 1986; Budd, 1998; Spratling, 2002; Spruston, 2008). This has led to theories of neuronal integration for layer 5 pyramidal cells that involve a “bottom-up” stream of information entering the basal dendrites and “top-down” signals coming to the apical tuft (Siegel, Körding and König, 2000; Larkum, 2013; Manita *et al.*, 2015). Moreover, it has recently been shown experimentally that when experiencing somatosensory stimulation, layer 5 pyramidal cells in S1 first exhibit an increase in firing rate corresponding to the bottom-up sensory input (ostensibly to the basal tree), and then, 30 ms later, receive top-down input to the apical tuft from M2 (Manita *et al.*, 2015). This indicates the presence of temporally segregated time windows in which the cell separately integrates input from the apical and basal tree. There is also work suggesting that plasticity rules may function differently in different regions of the cell (Gordon, Polsky and Schiller, 2006), again indicating that different regions of the cell might serve as input regions to distinct information pathways, and, as such, may have different priorities underlying the decision of when the cell will or will not fire. Taken together, the above studies strongly suggest that the apical tuft and basal dendrites can and should be studied as independent integration units.

### Inhibition

Our study made several simplifications in the learning and plasticity processes used as compared to those found in biology. Critically, our plasticity algorithm utilized only excitatory synapses and did not consider the effect of inhibition on learning. This is not because we believe that inhibition does not play a role in learning; on the contrary, inhibitory synapses are essential both for the learning process and in defining the input-output function of the cell (Wulff *et al.*, 2009; Kullmann *et al.*, 2012; Müllner, Wierenga and Bonhoeffer, 2015). However, by restricting ourselves to excitatory synapses, we were able to isolate important biophysical properties of excitatory synapses – namely the impact of synaptic saturation (the MESWs) that might have been masked in the presence of inhibition. Future work on the “biophysical perceptron” will include the role of inhibitory synapses; in this case special care must be taken to understand how inhibitory inputs interact with excitatory inputs on different locations of the cell (Gidon and Segev, 2012; Doron *et al.*, 2017). The addition of synaptic inhibition has the potential to increase the classification capacity of the cell (Julio Chapeton *et al.*, 2012), and localized inhibition may allow for additional forms of compartmentalized computation at the single neuron level.

The focus on excitatory synapses also enables our work to be directly compared to studies of excitatory perceptron-like learning done on Purkinje cells, such as the work of (Brunel *et al.*, 2004; Steuber *et al.*, 2007; Safaryan *et al.*, 2017) which have been classically conceived of as perceptrons (Marr, 1969; Albus, 1971). These studies demonstrated that detailed models of Purkinje cells can learn to discriminate between different patterns of input from the parallel fibers (PF) via a perceptron-like usage of long-term depression (LTD), which is known to occur in PF-Purkinje synapses. Crucially, the difference between the Purkinje cell’s responses to learned versus unlearned patterns was the duration of the pause between spikes in the Purkinje cell’s output subsequent to the presentation of PF input. Steuber et al. (Steuber *et al.*, 2007) argue that this pause duration-based learning depends on the mediation of calcium concentrations inside the cell. This is different from the more direct M&P-like mechanism, used in the present study, of synapses being weighted such that only certain input patterns will reach the cell’s spiking threshold.

### Nonlinearities and Alternative Plasticity Rules

Our focus on perceptron-like learning constitutes an additional simplification, as perceptron learning ignores the potential contributions that dendritic nonlinearities such as local NMDA spikes (Schiller *et al.*, 2000; Polsky, Mel and Schiller, 2004), dendritic Na^+^ spikes (Golding and Spruston, 1998; Sun *et al.*, 2014), dendritic Ca^2+^ spikes (Magee and Johnston, 1995; Kampa, Letzkus and Stuart, 2006; Cichon and Gan, 2015) may impact learning in classification tasks. Although a variety of dendritic nonlinearities are present in our L5 pyramidal cell model, we did not make explicit use of them in our plasticity rule. Indeed, some models of dendritic integration such as the Clusteron (Mel, 1991, 1992) and the two-layer model (Poirazi and Mel, 2001) treat the NMDA spike as critical for dendritic computation. In particular, these models treat clustering of nearby synapses, and “structural plasticity”, or the relocation of synaptic inputs within and between branches as crucial for learning (Trachtenberg *et al.*, 2002; Larkum and Nevian, 2008; Losonczy, Makara and Magee, 2008; Kastellakis *et al.*, 2015; Weber *et al.*, 2016; Mel, Schiller and Poirazi, 2017). The present study did not address the role of synaptic clustering in learning, a promising future direction would be to combine the weight-based learning rules used in our study with the structural plasticity algorithm as discussed in (Mel, 1992).

There are several other models of learning and plasticity that make use of neuronal biophysics and constitute promising opportunities for improving the learning ability of pyramidal cell models in a biologically plausible way. The calcium-based plasticity rule of (Graupner and Brunel, 2012) an exciting possibility for implementing perceptron-like learning in a more biological manner by making direct use of the experimentally observed mechanisms of plasticity in neurons. Because neurons exhibit some properties of multi-layered networks (Poirazi, Brannon and Mel, 2003; Beniaguev, Segev and London, 2019), it would also be valuable to explore more powerful learning algorithms that make use of the dendrites as a second (or higher) layer of computation as in (Schiess, Urbanczik and Senn, 2016). Alternatively, it may make sense to consider a different paradigm of dendritic learning, where the dendrites attempt to ‘predict’ the somatic output, allowing for forms of both supervised and unsupervised learning (Urbanczik and Senn, 2014).

### Timing

Another crucial element that remains to be studied in detailed biophysical models is the role of the timing of both the input and output of pyramidal cells in learning and computation. Regarding input timing, some theoretical work has been done on the M&P perceptron, which has been extended in a variety of ways to take into account several components of real neurons, like the Tempotron, which uses a leaky integrate and fire mechanism (Gütig and Sompolinsky, 2006) and can make use of conductance-based synapses (Gütig & Sompolinsky, 2009) to classify spatiotemporal input patterns. Regarding output timing and firing rate, learning rules like the one from (Gutig, 2016) can learn to solve the temporal credit-assignment by producing different spike rates for different inputs. Similarly, the Chronotron (Florian, 2012) considers learning rules that generate precisely timed output spikes. It is not clear to what extent these particular plasticity algorithms are truly “biological”, but there is no question that temporal sequence learning is an essential feature of the brain (Aslin, Saffran and Newport, 1998; Xu *et al.*, 2012; Moldwin, Schwartz and Sussman, 2017). The addition of a temporal dimension increases the classification capacity of the cell, as discussed in (Gütig and Sompolinsky, 2006).

### Broader Relevance

The present study shows that, by implementing the perceptron learning rule, layer 5 cortical pyramidal cells are powerful learning and generalization units, comparable – at the very least – to the abstract M&P perceptron. Other plasticity rules, which take into account synaptic clustering, input and output timing, and interaction between the apical and basal regions of pyramidal cells will be explored in further studies in detailed biophysical models in order to determine their biological plausibility and classification capacity. Until then, our study should be viewed as a baseline for comparison of any future work implementing learning algorithms in detailed biophysical models of neurons.

### Experimental Procedures

We utilized a detailed biophysical model of a layer 5 pyramidal cell written in NEURON with a Python wrapper (Carnevale and M.L. Hines, 1997; Hines, Davison and Muller, 2009). The parameters of the model, which includes numerous active mechanisms, are described in (Hay et al., 2011). Excitatory synapses were AMPA/NMDA-based synapses as in (Muller and Reimann, 2011) with synaptic depression and facilitation parameters set to 0. In both experiments, we placed all 1000 synapses in each pattern either on the soma, basal tree, or apical tuft according to a uniform spatial distribution.

### Memorization task

For the memorization task, each of the P patterns was generated by randomly choosing 200 out of the 1000 synapses to be activated. The patterns were then randomly assigned to either the positive or negative class. Patterns were presented to the cell by simultaneously stimulating the 200 active synapses with a single presynaptic spike at the beginning of the simulation. Simulations of the neuron were run with a Δt of 0.1 ms for a total of 100 ms. Patterns were considered to have been classified as “positive” if they produced at least one spike within the 100 ms time window and as “negative” if no spikes occurred.

We utilized an “online” version of the perceptron learning algorithm, applying the plasticity rule every time a pattern was presented to the neuron. Also, because we limited our analysis to excitatory synapses, we use the modified algorithm proposed in (Amit, Wong and Campbell, 1999) for sign-constrained synapses, which ensures that synaptic weights never become negative.

The algorithm works as follows: An presynaptic input pattern ***x*** is presented to the neuron, where ***x*** is a vector consisting of 1000 binary inputs, each of which is labeled *x*_*i*_ and associated with a particular synapse on the dendritic tree with synaptic weight *w*_*i*_ (for conductance synapses, this is the excitatory conductance of the synapse, 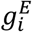). Each pattern has a target value, *y*_0_ ∈ {1, −1}, associated with it, where 1 means “should spike” and −1 means “shouldn’t spike.” When the pattern is presented to the neuron via simultaneous activation of all the synapses in the pattern, the soma of the neuron will produce a voltage response. If that voltage response contains at least one spike within 100 ms, we set the output variable *y* = 1. If the voltage response does not contain any spikes, we set *y* = −1. For each presynaptic input pattern, the plasticity rule for synapse *i* to update its weight *w*_*i*_ at time is defined as:

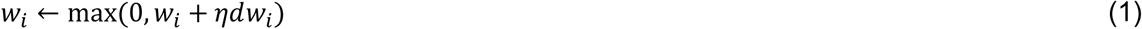

Where *dw*_*i*_ is defined as:

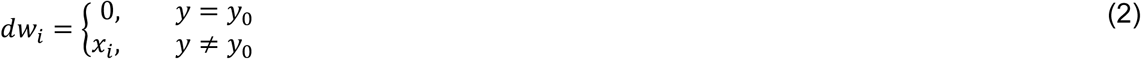

and *η* is the learning rate.

In other words, if the target output is the same as the actual output of the neuron, we do nothing. If the target is “should spike” and the neuron does not spike, we increase the weight of all synaptic inputs that were active in the pattern. If the target is “shouldn’t spike” and the neuron does spike, we decrease the synaptic weights of all synaptic inputs that were active in the pattern, unless that would decrease the synaptic weight below 0, in which case we reduced the weight of that synapse to 0.

The accuracy of the neuron’s output was calculated after each epoch, which consisted of a full pass of presenting each pattern (in random order) to the neuron. To ensure that accuracy improved on every epoch and reached a reasonable asymptote for all conditions, we set the learning rate *η* to 0.002 for the condition with AMPA/NMDA conductance synapses and an active tree, and a rate of 0.19 for the condition with current synapses. We also used the “momentum” technique (Rumelhart, Hinton and Williams, 1986) to improve learning speed. An average simulation time for a complete run of the learning algorithm for the memorization task (i.e. 100 epochs) was several days. Results shown in figures 2A-2D and 3D are averaged over 10 runs of the memorization task.

### M&P model (not constrained by synaptic battery)

To compare the BP to an equivalent M&P perceptron (Figures 2A-B and 5A-B) we used a M&P perceptron with only excitatory weights as described in (Amit, Wong and Campbell, 1999) (See Equation (1) and (2) above). A M&P neuron with no inputs would have a “bias” input value of - 77.13 to mimic the resting potential of the BP and a “spiking threshold” of −22.8 to mimic the voltage spiking threshold of the biophysical neuron. The learning algorithm used a learning rate *η* of 0.0008 (hand-tuned to optimize performance) which was dynamically modified in the learning algorithm via the momentum technique (Rumelhart, Hinton and Williams, 1986).

### MESW calculation and MESW-constrained M&P model

To calculate the MESWs for the L5PC model, we voltage clamped each dendritic segment in the neuron model to the synaptic reversal potential of 0. The MESW for a dendritic segment is defined as the difference between the somatic resting potential and the peak depolarization obtained at the soma within 100 ms after synaptic activation. (Figure 3A).

To create an MESW-constrained M&P model for the apical tuft, we calculated the distribution of MESWs per unit length of the dendritic membrane in the apical tuft. (The median and quartile values of the MESWs for all synaptic placement conditions are shown in the box-and-whisker plot in Figure 3C.) We then created an M&P neuron where each weight was individually given a “cap” drawn randomly from the apical tuft MESW probability distribution which would prevent the weight of that input from increasing above a certain value. In other words, if the plasticity algorithm (Equation 1) would bring *w*_*i*_ to be greater than cap, *c*_*i*_, we would “freeze” the weight at *c*_*i*_. Formally, this means that the plasticity rule in the case of an error for the MESW-capped neuron is

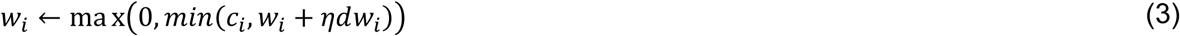

Where *η* and *dw*_*i*_ are as defined above in Equation (2).

### Generalization task

In the second task (generalization), we created two underlying patterns of 1000 synapses each, where 200 synapses were active, as in the memorization task. These patterns were then corrupted by flipping a given number synapses (0, 100, or 200, depending on the condition) and presented to the neuron. To maintain the sparsity of the patterns, half of the flipped bits were switched from active to inactive and the other half switched from inactive to active. For example, in the condition with 100 flipped bits, 50 out of the 200 previously active synaptic inputs were flipped to inactive, and 50 out of the 800 previously inactive synaptic inputs were switched to active.

In every epoch of the learning task, we presented the neuron with 50 noisy patterns generated by the first underlying pattern and 50 noisy patterns generated by the second underlying pattern for a total of 100 patterns per epoch (the order of the presentation of patterns from the two underlying patterns was also randomized). We set the learning rate *η* to 0.25 for the condition with AMPA/NMDA conductance synapses and an active tree, and a rate of 10 for the condition with current synapses. Similar to the memorization task, we used the online perceptron learning rule with the momentum modifier. In this task we only ran the algorithm for 5 epochs, as this was enough for the learning to achieve a plateau. Results shown in Figure 5A-5D are averaged over 25 repetitions of the generalization task.

Simulations were all performed using Neuron v.7.6 (Carnevale and M.L. Hines, 1997; Hines, Davison and Muller, 2009) running on a multi-core cluster computer. An average simulation time for a complete run of the learning algorithm for the generalization task (i.e. 5 epochs) was several minutes.

## Acknowledgements

We would like to thank Oren Amsalem for his assistance in many aspects of this work, particularly his help with the use of the NEURON software. We also would like to thank David Beniaguev, Guy Eyal, and Michael Doron for their insightful discussions about machine learning and neuronal biophysics. Itamar Landau provided several useful comments to an early version of this work, inspiring important revisions. Additionally, we appreciate Nizar Abed’s support in maintaining the computer systems used to perform our simulations and data analysis. This work was supported by the Drahi family foundation, the EU Horizon 2020 program (720270, Human Brain Project), a grant from Huawei Technologies Co., Ltd., and by a grant from the Gatsby Charitable Foundation.

## Notes

#### Summary of Updates

Minor edits for formatting and style, additional references in discussion

